# The many nuanced evolutionary consequences of duplicated genes

**DOI:** 10.1101/366971

**Authors:** Ashley I. Teufel, Mackenzie M. Johnson, Jon M. Laurent, Aashiq H. Kachroo, Edward M. Marcotte, Claus O. Wilke

**Author notes:** Present address: Institute for Systems Genetics, Department of Biochemistry and Molecular Pharmacology, New York University Langone Health, New York, NY, USA. Present address: The Department of Biology, Centre for Applied Synthetic Biology, Concordia University, Montreal, QC, Canada.

## Abstract

Gene duplication is seen as a major source of structural and functional divergence in genome evolution. Under the conventional models of sub- or neofunctionalizaton, functional changes arise in one of the duplicates after duplication. However, we suggest here that the presence of a duplicated gene can result in functional changes to its interacting partners. We explore this hypothesis by *in-silico* evolution of a heterodimer when one member of the interacting pair is duplicated. We examine how a range of selection pressures and protein structures leads to differential patterns of evolutionary divergence. We find that a surprising number of distinct evolutionary trajectories can be observed even in a simple three member system. Further, we observe that selection to correct dosage imbalance can affect the evolution of the initial function in several unexpected ways. For example, if a duplicate is under selective pressure to avoid binding its original binding partner, this can lead to changes in the binding interface of a non-duplicated interacting partner to exclude the duplicate. Hence, independent of the fate of the duplicate, its presence can impact how the original function operates. Additionally, we introduce a conceptual framework to describe how interacting partners cope with dosage imbalance after duplication. Contextualizing our results within this framework reveals that the evolutionary path taken by a duplicate’s interacting partners is highly stochastic in nature. Consequently, the fate of duplicate genes may not only be controlled by their own ability to accumulate mutations but also by how interacting partners cope with them.

## Introduction

Gene duplication is a major driver of functional divergence (Lynch and Conery, 2000; Ohno, 1970). Duplicate genes provide an additional source of genetic material that is free from the selective constraints experienced by the original gene copy (Ohno, 1970). While freedom from these selective constraints often results in the duplicate losing function and being expelled from the genome, some duplicate genes acquire functional changes resulting in their preservation. The long term preservation of a duplicated gene is often attributed to one of two mechanisms. Under the first, known as *neofunctionalization*, the duplicate gene acquires a novel and beneficial functional change leading to the gene’s retention. Under the second, called *subfunctionalization*, both the original and the duplicate gene copies acquire partial loss-of-function mutations, requiring their mutual retention to perform the original function (Dittmar and Liberles, 2011; Innan and Kondrashov, 2010). While the preservation of duplicate genes is rare, it is thought to be more likely to occur if a newly duplicated gene remains in the genome for an extended period of time. This extended window of time allows for an increased opportunity for mutational and selective forces to reshape the function of the duplicate gene (Konrad et al., 2011). Further, these mechanisms are not mutually exclusive. The initial retention of a duplicate could be due to subfunctionalization that eventually leads to neofunctionalization (Roth et al., 2007). When multiple genes are duplicated, selective pressure to maintain dosage balance can extend the initial retention period and serve as mechanism for later sub- or neofunctionalization (Teufel et al., 2016). Genome architecture may also play a role in the initial retention of a duplicate, as it may be more difficult to expel duplicates depending on their location.

While a number of theories offer explanations of how specific forces foster the functional diversification of duplicate genes, little attention has been given to the secondary effects duplicates have on the evolution of their interacting partners. Considering that the rate of gene duplication can be similar to or exceed the rate of synonymous substitutions (Lipinski et al., 2011), the ability of duplicated genes to affect the evolutionary trajectory of interacting partners may be significant. We hypothesize that the traditional view, under which functional changes occur within the duplicate itself, may not capture the full spectrum of evolutionary outcomes for a system of interacting genes after a duplication.

Here we explore the evolution of a heterodimeric protein complex when one subunit is duplicated. Even in this simple system of three interacting proteins, there are a number of ways that the proteins could be impacted by the presence of a duplicate. To examine the evolutionary consequences of gene duplication, we simulate protein evolution and assess the stability of its subunits, their ability to bind, and the mechanisms driving functional change. We do so under various selection scenarios for several heterodimeric structures. We find that the presence of a duplicate gene can influence the evolution of each member in a proteinprotein interaction. Additionally, we observe a surprising amount of variability in how protein-interaction networks cope with dosage imbalance, and we introduce a framework for describing the impact imbalance can have on interacting partners. Our results highlight that the presence of a duplicate gene can affect the evolution of each of its interacting partners and that this impact depends both on protein structure and stochastic events.

## Results

We examine the evolution of a simplified protein-interaction network after a partial duplication. While the traditional view of gene duplication predicts that the duplicate gene will escape selective pressure, we suggest that the presence of a duplicate can impose a new set of selective constraints on its interacting partners. Considering a relatively simple system of a heterodimer SUMO ubiquitin-like protein complex (PDB ID: 2EKE, Duda et al. 2007) whose two subunits we refer to as A and B, we impose a duplication event that results in a redundant copy of the B subunit (denoted as B’) (Fig. 1).

**Figure 1:**
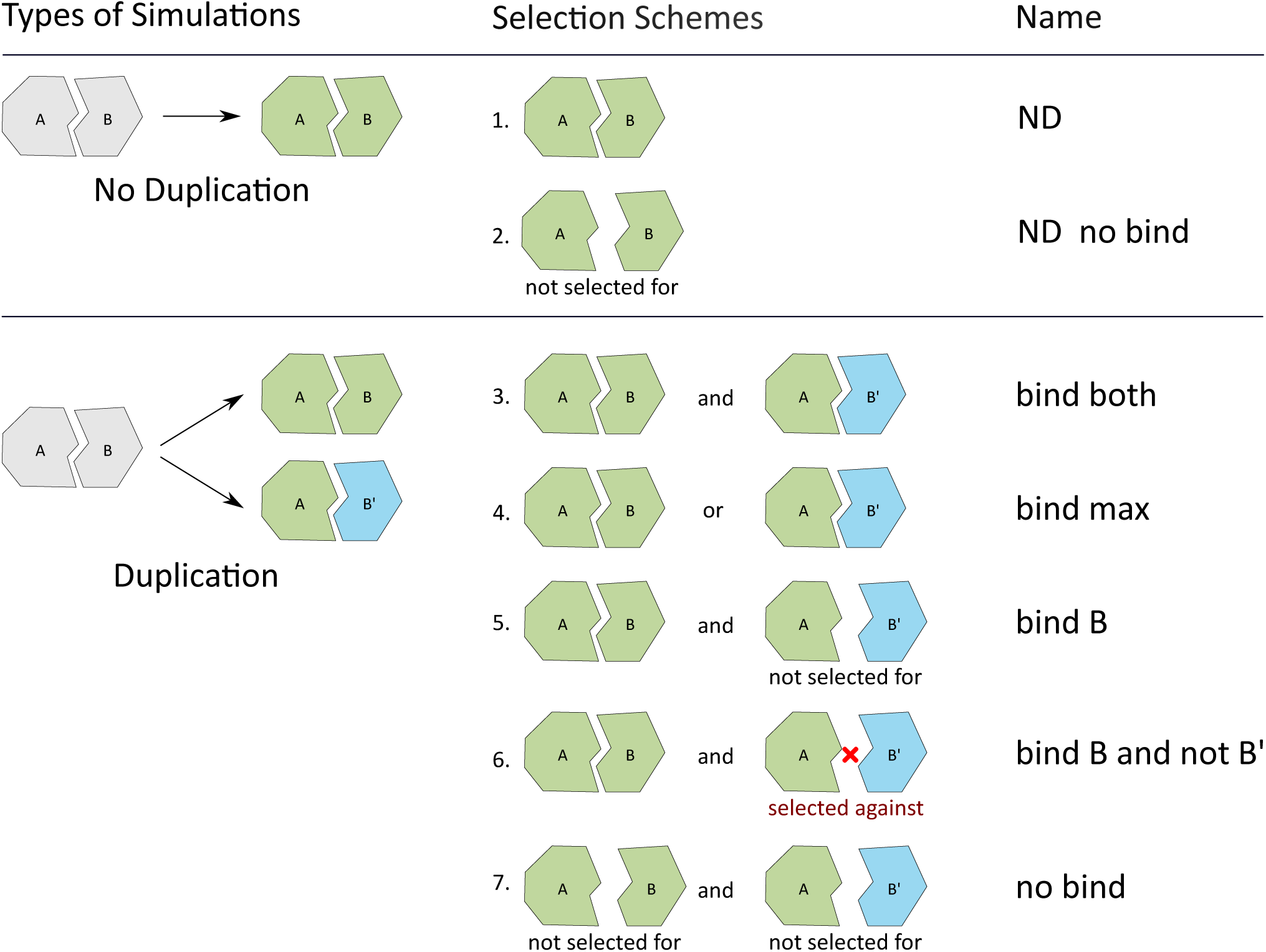
Selection schemes. We simulate evolution under seven different selection schemes, five of which include the duplication of protein B, denoted as B’. All simulations assume selection for the stability of A, B, and B’ if applicable. Two simulations assume that a duplication event does not occur. Selection scheme 1 (no duplication, ND) describes a scenario without duplication where the stability of the A–B interface is included in the fitness function. Selection scheme 2 (ND no bind) describes a scenario without duplication where the stability of the A–B interface is not included in the fitness function. To examine how the duplication of a subunit affects evolutionary dynamics, we consider five additional selection schemes. Selection scheme 3 (bind both) describes a scenario where both duplicates (B and B’) need to bind A, and the stability of both the A–B and the A–B’ interface is included in the fitness function. This type of selection pressure could occur in a situation where increased dosage of B is beneficial. Selection scheme 4 (bind max) describes a competition scenario where the stability of only one interface, that of the maximum stability of binding for either the A–B or the A–B’ interface, is included in the fitness function. Selection scheme 5 (bind B) describes the process of B’ nonfunctionalization, and the stability of only the A–B interface is included in the fitness function. Selection scheme 6 (bind B and not B’) describes diversifying selection, and the stability of the A–B interface is included in the fitness function while the stability of the A–B’ interface is used as a fitness penalty. This sort of selection scenario mimics that of dosage imbalance, where an excessive amount of unbound B’ is harmful. Selection scheme 7 (no bind) describes a control duplication experiment where the stability of neither the A–B nor the A–B’ interface is included in the fitness function.

To examine how protein-interaction networks are impacted by duplication, we implement evolutionary simulations under seven different selection schemes (Fig. 1). These selection schemes are chosen to reflect different evolutionary scenarios that have been hypothesized to contribute to duplicate gene divergence. Two of these experiments are control simulations that do not include a duplication event (Fig. 1, selection schemes 1 and 2). The five remaining simulations include a duplicate of B, referred to as B’ (Fig. 1, selection schemes 3 through 7). To examine the influence of protein structure, a subset of these experiments (Fig. 1, selection schemes 3, 5, and 6) are repeated assuming that A is the duplicate protein (denoted as A’) and with the heterodimer protein structure of antifungal protein KP6 (PDB ID: 4GVB, Allen et al. 2013).

## Stability of protein interfaces and structures

To assess how the presence of a duplicate gene influences the evolution of protein interfaces, we measure the stability of the A–B interface over the course of the simulation (Fig. 2). The resulting interface stability can be described by three general dynamic patterns. Selection schemes that do not select for binding (selection schemes 2 and 7) result in a destabilization of the protein interface. Most selection schemes that do select for binding (selection schemes 1 and 3-5) consistently maintain interface stability. However, selection scheme 6, which corresponds to deleterious dosage imbalance, causes initial destabilization of the interface followed by a recovery of stability (Fig. 2, dark blue line).

**Figure 2:**
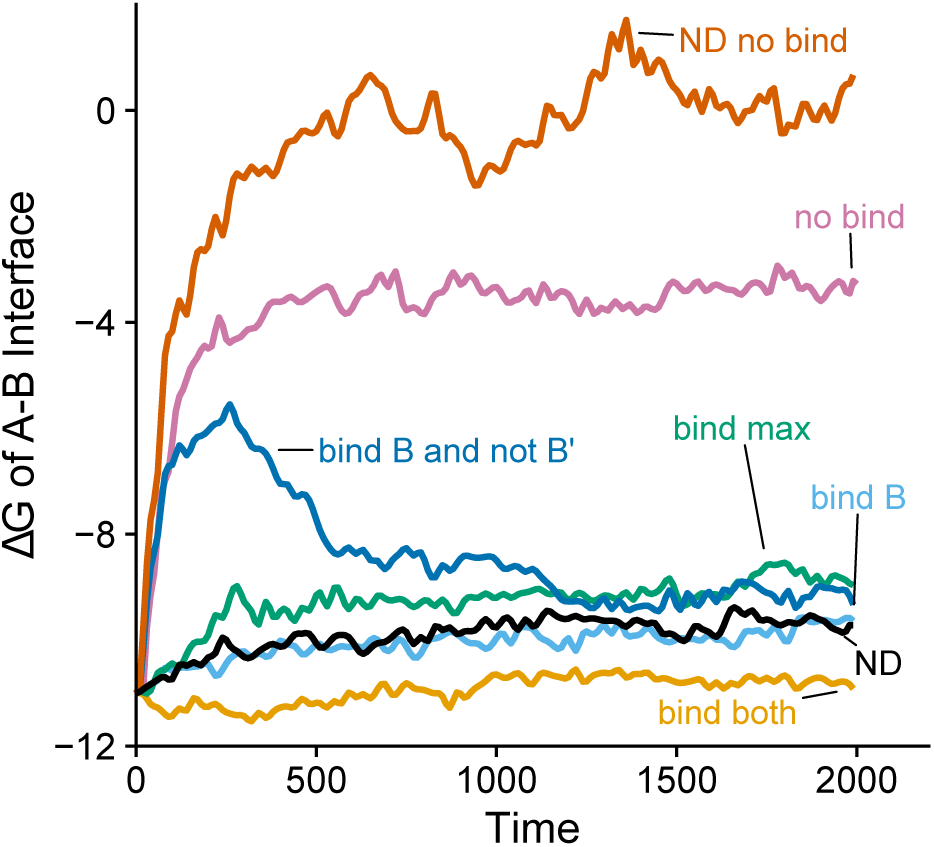
Stability of the A–B interface versus time, for all 7 selection schemes of Fig. 1. Δ*G* values are averaged over replicate simulations. Under selection schemes that do not reward binding, the interface stability tends to decay rapidly (more positive Δ*G* indicates less stable binding). By contrast, selection schemes that do reward binding tend to maintain the interface stability throughout. One exception to this pattern is the bind B and not B’ case, which first shows rapid destabilization of the A–B interface, followed by subsequent regaining of binding stability.

While the dynamics of selection schemes 1-5 and 7 are expected, the destabilization of the A–B interface under deleterious dosage imbalance (selection scheme 6) is unexpected and surprising. This observation demonstrates that selection to avoid binding B’ can result in a change to the A–B interface to avoid A–B’ binding. Hence, selection for a functional change of B’ impacts how the A–B interface evolves, forcing a temporary reduction in the stability of the A–B interface. We also examine the stability of the A–B’ interface and find that it displays similar dynamics to the stability of the A–B interface under most of the selection scenarios (Fig. S1). However, under the selection scenario to avoid binding B’, the A–B’ interface quickly destabilizes (Fig. S1, dark blue line), as one would expect.

We measure the stability of the duplicate protein, B’, to assess if any of our selection schemes have other effects on the evolving protein (Fig. S2). We find that most of our selection scenarios (selection schemes 3-5, 7) result in a consistent level of stability. However, under deleterious dosage imbalance (selection scheme 6), we find that B’ is destabilized early on in the simulation (Fig. S2, dark blue line). It appears that B’ is never able to fully recover from this initial destabilization. These findings demonstrate that selection for diversifying functionality, such as the loss of binding ability, may initially cause a destabilization of a protein’s structure. After functional changes occur, the stability of the structure may then be refined. We obtain similar results for interface and duplicate stability when we duplicate A instead of B (Figs. S3, S4) and when we simulate using the antifungal protein (Figs. S5, S6).

## Retention of ancestral function

To further examine how the presence of a duplicate affects the A–B binding interface, we compare the functionality of the evolved interface to its ancestral function. To compare our evolved and ancestral functions, we assess if our evolved A protein can functionally bind the ancestral B protein. We consider binding to be non-functional if the thermodynamic stability of binding does not exceed a protein-specific minimal threshold (see Methods). The percentage of simulations where A retains its ability to bind the ancestral B is given in Fig. S7A for each selection scenario. Generally, it appears that selection for A–B binding, as implemented in selection schemes 3-5, results in a conservation of ancestral function of the A protein, consistent with prior work in a non-duplicated context (Kachroo et al., 2015).

However, when selection acts to avoid binding the duplicate B’ (selection scheme 6), there is a sharp decrease in the ability of A to perform its ancestral function (S7A, dark blue line). This indicates that the A subunit changes substantially in order to avoid binding B’, such that only around 20% of the A proteins are able to bind the ancestral B by the end of the simulation. This result shows that a functional change can occur in a non-duplicated interacting partner in response to the presence of a duplicate.

We also evaluate how our duplicated proteins, B and B’, functionally diverge based on their ability to bind to the ancestral A protein (Fig. S7B, C). The ability of the B (Fig. S7B) and B’ (Fig. S7C) proteins to bind the ancestral A protein decreases more sharply than observed for the A protein. This suggests that the duplicated proteins diverge in function more so than the A protein. Notably, both duplicates similarly retain the ability to bind the ancestral A protein under most selection scenarios, with the exception of selection to avoid binding B’ (Fig. S7C, dark blue line).

### Pathways of adaptation to dosage imbalance

Considering the destabilization of the A–B interface and of the B’ structure observed when dosage imbalance is selected against (Figs. 2, S2, dark blue lines), as well as the evidence that the A subunit appears to undergo functional change (Fig. S7A, dark blue line), we assess how deleterious binding is escaped on a biochemical level. We use a chi-square test to determine which sites have differing amino acid compositions between the bind both (selection scheme 3) and bind B and not B’ (selection scheme 6) experiments at every 10th substitution for the first 250 substitutions.

We find that the amino acid compositions of site 20 in A and 46 in B’ differ significantly (*α* = 0.05, Benjamini and Hochberg 1995 corrected). The number of generations before these sites display significant differences in their amino acid composition also differs. Site 20 in A displays a significant difference by generation 70, while site 46 in B’ displays a significant difference by generation 80. This difference in timing suggests that diversifying changes first occur in A and then in the duplicate B’. Further, the loss of A–B’ binding occurs via modifications to site 20 in A and 46 in B’.

To investigate if other downstream effects occur after functional change, we repeat the same chi-square analysis at every 100th substitution. Other sites in protein B’ (9, 44, 47, and 68) also display differing amino acid compositions between the two experiments, though these sites do not differ significantly until after 350 generations. These changes are most likely secondary effects that occur after A has escaped binding to B’. Notably, each of the sites that differ in their amino acid compositions are located in the binding interface (Fig. 3).

**Figure 3:**
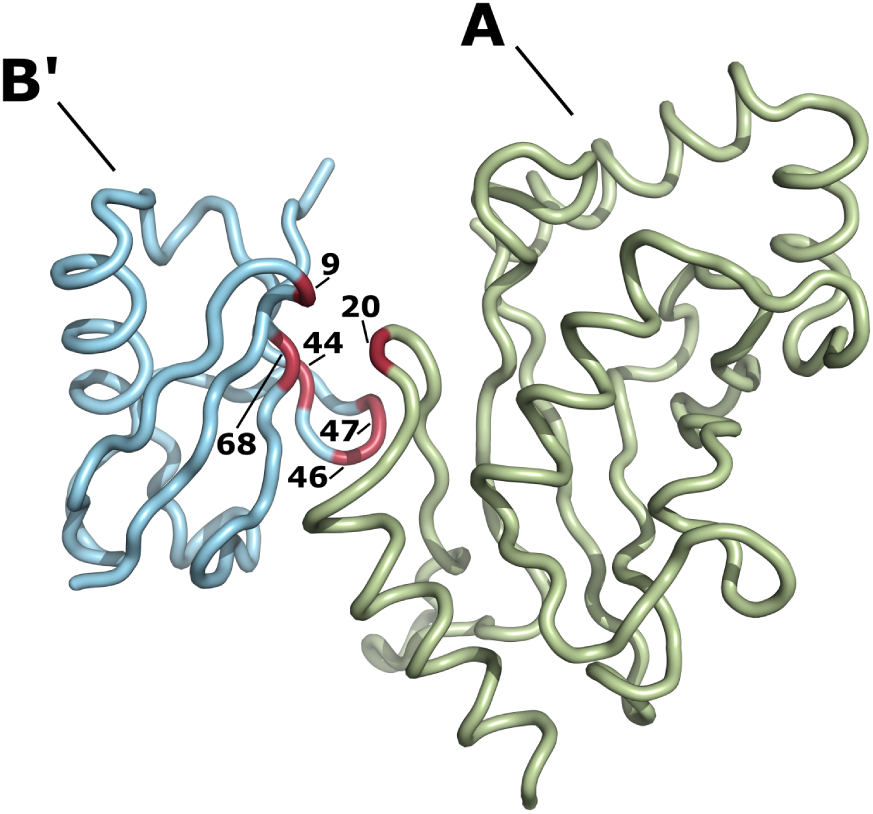
Significantly differing sites after adaptation to different selection schemes. The highlighted sites differ significantly in their amino acid composition between the bind both and the bind B and not B’ simulations. These sites include site 20 in protein A and sites 9, 44, 46, 47, and 68 in protein B’. No sites were found to significantly differ in their amino acid composition in protein B.

To examine the properties of sites that significantly differ in their amino acid composition, we assess how residues at these sites contribute to binding using a “stickiness” scale (Levy et al., 2012), which describes the propensity of amino acids to be in protein-protein interfaces. The distributions of amino acid stickiness of site 20 in A and site 46 in B’ is shown for the first 250 substitutions (Fig. S11). Under the bind both selection scheme, the relative stickiness of both of these residues is maintained across time, indicating that maintaining the initial distribution of stickiness is important for the retention of binding. Further, it appears that site 20 initially becomes less sticky to avoid binding B’, suggesting that increasing the propensity for amino acids that are less likely to be involved in protein-protein interactions is crucial to avoid B’ binding. Site 46 in B’ (Fig. S11B) appears to increase its propensity for interface residues while under selection to avoid binding B’, suggesting that perturbing the original composition of site stickiness is crucial to avoid B’ binding.

Additionally, we examine the long-term dynamics of sites with differing amino acid compositions at every 100th generation, and observe similar dynamics in terms of the stickiness (Fig. 4) and mass of these residues (Fig. S12). Most notably, while site 20 in protein A initially decreases in its stickiness (Fig. 4A) and mass (Fig. S12), the site later regains some of its stickiness and mass. Other sites in the B’ subunit also display differing distributions in terms of their stickiness (Fig. 4). The persistence of differing amino acid compositions at sites in both the non-duplicated interacting partner and the deleterious duplicate throughout the simulation suggests that both the A and the B’ subunit undergo diversifying changes.

**Figure 4:**
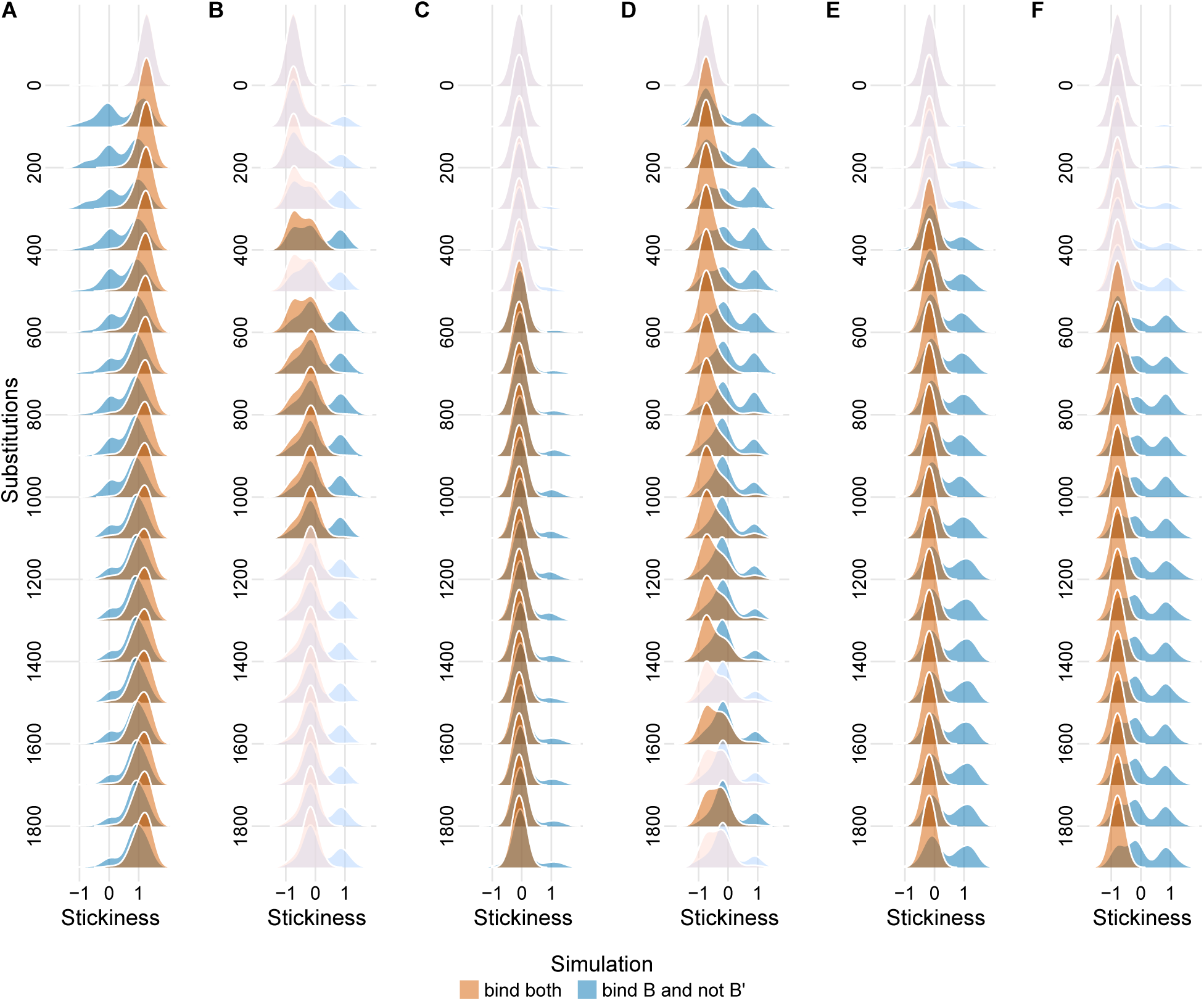
Distribution of stickiness for sites with differing amino acid compositions, shown at every 100th generation. Grayed-out distributions indicate time points at which sites do not display significant differences. (A) Site 20 in protein A. While the stickiness of site 20 in protein A in the bind B and not B’ scenario is initially reduced, some stickiness is regained and by the end of the simulation the distributions of residue stickiness are similar for the two selection schemes. This observation suggests that once dosage imbalance is escaped, the A–B interface is refined and reoptimized. (B) Site 9 in protein B’. This site displays a slight shift toward more sticky residues under selection to avoid binding B’, though a larger portion of this distribution still resembles the bind both scenario. However, distributions differ significantly only from generation 400 to generation 1100, and this transient behavior may be related to the re-stabilization of the B’ structure. (C) Site 44 in protein B’. For this site, the differing amino acid composition between the two selection schemes does not appear to be reflected in the distribution of amino acid stickiness. (D) Site 46 in protein B’. The dynamics at this site suggest that selection to avoid binding B’ initially shifts the distribution towards stickier residues. Interestingly, this shift also occurs under the bind both selection scheme, just later in time. It appears that selection to avoid binding B’ results in an accelerated exploration of sequence space. (E) Site 47 in protein B’. (F) Site 68 in protein B’. Sites 47 and 68 increase in stickiness under selection to avoid binding B’. However, this effect sets in only around generation 500, indicating that these changes are related to the re-stabilization of B’.

To investigate how deleterious binding is lost when A’ is the duplicate subunit, we again measure the stickiness (Fig. S13) and mass (Fig. S14) of residues found to have significantly different distributions of amino acids across time. However, only sites in A’ appear to have continuous differences across time in their amino acids compositions (Fig. S13A, C), indicating that diversifying changes occur in the A’ subunit. When examining the stickiness (Fig. S15) and mass (Fig. S16) of sites that significantly differ in simulations of the antifungal protein complex we find that sites in both B and B’ significantly differ across time. Interestingly, the presences of the deleterious duplicate B’ appears to influence the mass of residues at a site in B (Fig. S16B), despite the fact that these two proteins do not directly interact. We examine why changes towards smaller amino acids at this site occur by substituting a glycine into this position of the protein structure used to initialize these simulations. We find that this substitution increases the A–B binding stability (ΔΔG = −1.06); however, it decreases the stability of the B protein (ΔΔG = 5.48). Hence, changes to this site appear to be a compensatory mechanism to stabilize the A–B interface during selection to avoid binding B’, but at the cost of structural destabilization. Upon inspection of the stability of B, we do indeed observe a slight destabilization early on in the simulation (Fig. S17, blue line).

We further examine how these functionally diversifying changes occur by tracking the location of the first 250 substitutions relative to the binding interface. We find that selection to avoid binding B’ results in a preference for changes to interface residues in the A and B’ proteins beyond what is observed in the bind both selection scheme (Fig. S8). We obtain similar results when we duplicate A instead of B (Fig. S9) and when we simulate using the antifungal protein structure (Fig. S10). The combination of these results, along with our analysis of critical interface sites, demonstrates that deleterious dosage imbalance results in an increased rate of interface substitutions as well as changes to the amino acid composition of critical interface sites.

As each simulation of deleterious dosage imbalance (selection scheme 6) initialized with a different structure results in diversifying changes in different subunits, we conclude that the mechanisms of functional differentiation are dependent on a protein’s structure. In our systems of three interacting proteins we have observed three different outcomes, though five other combinations are theoretically possible as well (Fig. 5). In our initial simulation, we have found that functional changes occur in the duplicate itself and in the non-duplicated interacting partner (option 5 in Fig. 5). When duplicating the A protein, we find that the changes essential for escaping deleterious binding occur in the duplicate itself (option 1 in Fig. 5). Finally, in the simulations initialized with the antifungal structure, we observe that changes occur in the interfaces of both of the duplicated proteins (option 4 in Fig. 5). Notably, in each of our simulations the deleterious duplicate displays functional changes.

**Figure 5:**
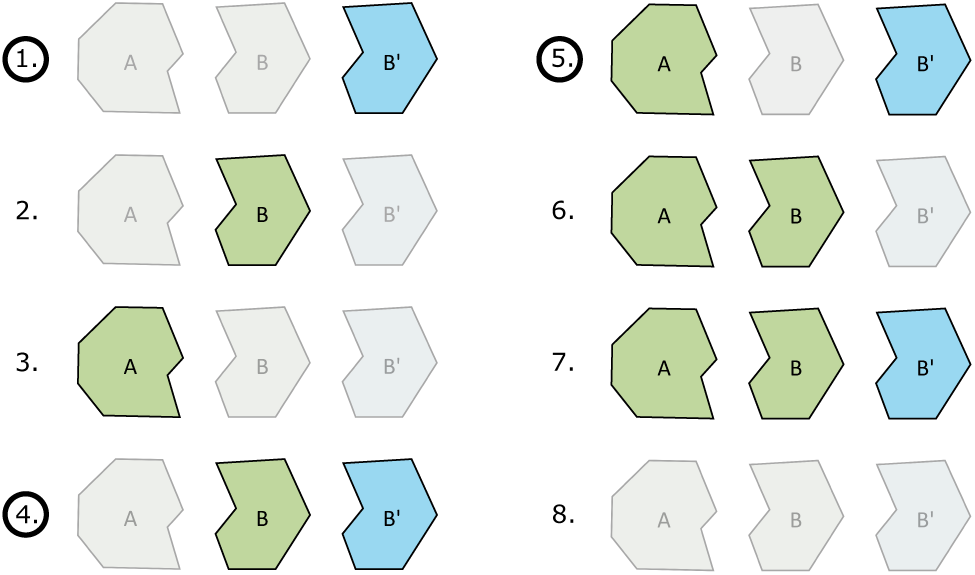
All possible ways in which the proteins in a three-member network could adapt after duplication. Grayed-out structures indicate no adaptation and structures in full color indicate diversifying change. In options 1 and 2, only one of the duplicated interacting partners acquires diversifying changes. Option 3 describes a situation where only the non-duplicated interacting partner accumulates diversifying changes. Option 4 describes the case where both of the duplicated genes acquire diversifying changes. In options 5 and 6, the non-duplicated partner and one of the duplicate partners accumulate diversifying changes. Finally, in option 7 all proteins and in option 8 none of the proteins acquire diversifying changes. Black circles denote options that we observe in this study.

### Beyond sub- and neo-functionalization: The impact of functional change on interacting partners

To quantify the impact that deleterious dosage imbalance has on its interacting parters, we examine how the interacting partners functionally diversify in response to the duplicate’s presence. Functional change is assessed by measuring an evolved protein’s ability to bind extant and ancestral partners (see Methods for details). To describe how a deleterious duplicate’s interacting partners cope with its presence, we define functionalization pathways in terms of a protein’s ability to bind extant and ancestral partners (Table 1). When a protein loses the ability to bind its current partner and does not later regain this ability, we refer to this process as *defunctionalization*. When a protein is found to consistently bind its current interacting partner throughout the simulation, but loses the ability to bind its ancestral partner at any point, we refer to this process as *isofunctionalization*. When a protein loses the ability to bind its current partner and then regains the ability later, we refer to this process as *refunctionalization*. After refunctionalization, three distinct fates are possible. If binding to the ancestral partner is permanently lost after refunctionalization, we refer to this process as *hard isofunctionalization*. If binding to the ancestral partner is regained at any point after refunctionalization, we refer to this process as *soft isofunctionalization*. Finally, if binding to the ancestral partner is retained across evolution after refunctionalization, we refer to this process as a *functional reacquisition*.

For the case of selection to avoid binding B’ (selective scheme 6), we compare the fractions of simulations resulting in each functionalization pathway from the perspective of the non-duplicated interacting partner (Fig. 6A) and the non-deleterious duplicate (Fig. 6B), for simulations initialized with different structures. Notably, each structure results in distinct fractions of functionalization pathways, suggesting that protein structure plays a role in how functional change is achieved. Further, the variation of diversification mechanisms displayed by each protein structure suggest that stochastic events shape how functional change is achieved in replicate simulations.

**Figure 6:**
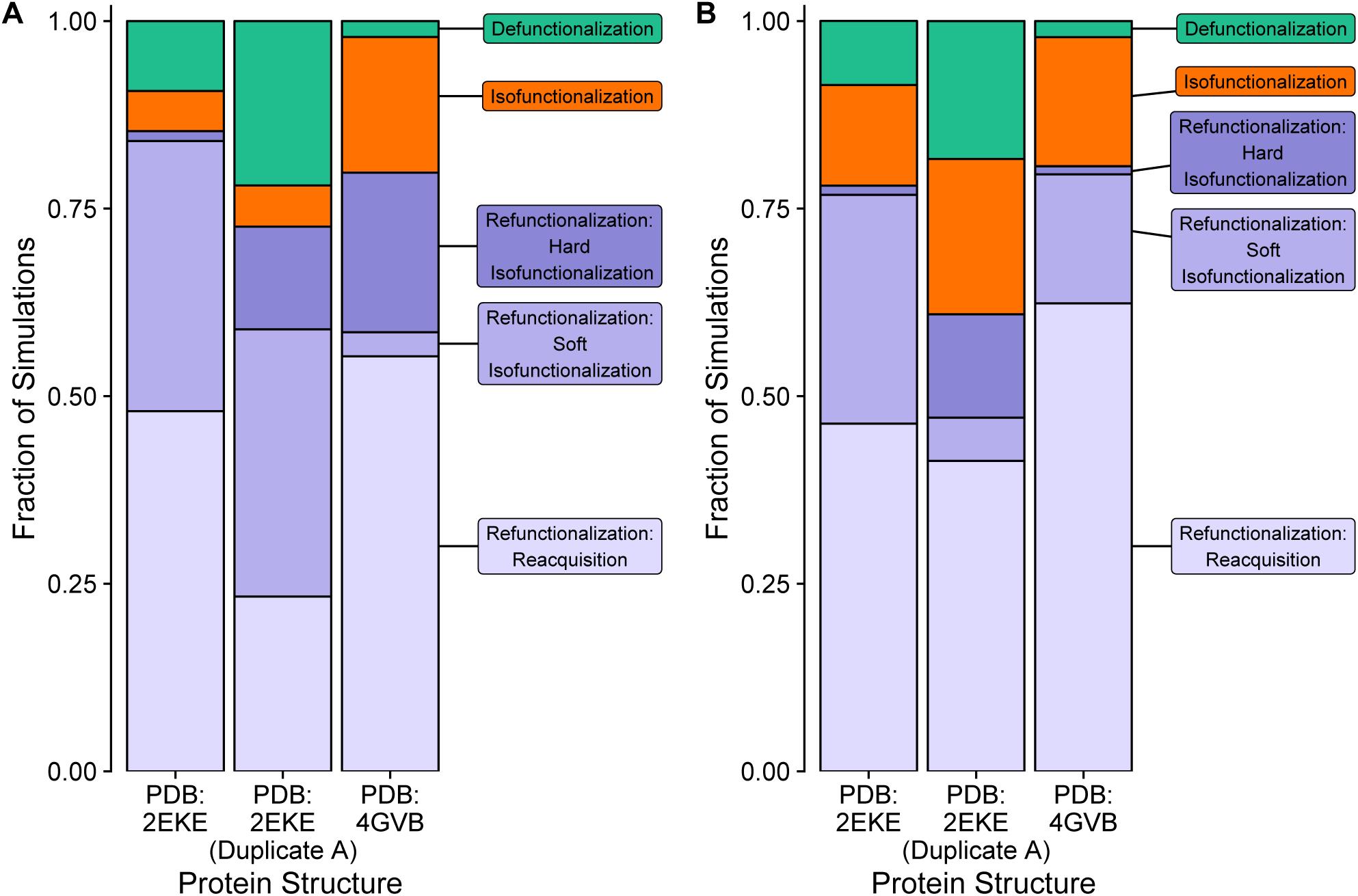
Comparison of functionalization pathways under dosage imbalance. Each bar represents an experiment initialized with a different starting structure. (A) Fraction of functionalization pathways of the non-duplicated interacting partner. (B) Fraction of functionalization pathways of the duplicated interacting partner that is under selection to maintain binding. Data is not shown for the deleterious duplicate because it defunctionalizes in all replicates.

For the non-duplicated interacting partner (Fig. 6A), only a small fraction of our simulations escape dosage imbalance without the temporary loss of the ability to bind the extant partner (isofunctionalization). This indicates that avoiding binding a deleterious duplicate while maintaining binding to another duplicate is possible, though this outcome is relatively rare. Most of the non-duplicated interacting partners undergo refunctionalization, a process that reflects the destabilization of the interface observed early on in each of the simulations (Figs. 2, S3, S5). After refunctionalization occurs, the non-duplicated partner’s interface can be reshaped in several different ways. In the case of the simulations initialized with the ubiquitin-like protein complex (PDB: 2EKE) and the antifungal protein (PDB: 4GVB), this reshaping often results in a protein interface that is still able to perform the ancestral function (reacquisition), though this is observed less often when A is the duplicated subunit (Fig. 6A). In each of these simulations, we also observe a notable fraction of cases where binding to the ancestral partner is entirely or intermittently lost after refunctionalization. This result indicates that, in a substantial portion of our simulations, the interface of the non-duplicated interacting partner functionally diversifies from that of the ancestral interface. This finding reinforces the idea that the presence of a deleterious duplicate can result in lasting functional change of the non-duplicated partner.

Examining the functionalization pathways of the non-deleterious duplicate (Fig. 6B), it appears that a larger fraction of non-deleterious duplicates undergoes isofunctionalization than does their non-duplicated partner, implying that functional maintenance is more prominent for the non-deleterious duplicate. We also observe that most of the non-deleterious duplicates undergo a phase of refunctionalization; however, this pathway often results in reacquisition of the ancestral function. Notably, in only a few simulations do we see permanent loss of the ability to perform the ancestral function after refunctionalization (hard isofunctionalization), while a few waver on this ability (soft isofunctionalization). A comparison of the non-duplicate and non-deleterious duplicate fractions of functionalization pathways (Fig. 6A, B) suggests that the effect of deleterious dosage imbalance affects the non-duplicated interacting partner more substantially than it does the non-deleterious duplicate.

## Discussion

We have examined how protein-interaction networks evolve when one member of an interaction network is duplicated. Under a number of different selection schemes, we find that how selection acts on duplicates can impact how their interacting partners evolve. Most notably, we find that escaping deleterious dosage imbalance can be achieved through several mechanisms, summarized in Fig. 5, though five other mechanisms we have not observed are also theoretically possible. Considering that a different combination of changes to interacting partners is observed under each type of simulation instantiated with a different structure, this finding suggests that protein structure plays a substantial role in the location of diversifying changes. Interestingly, in two of our experiments, we find that changes to the original binding interface occur due to the presence of the deleterious duplicate. This functional modification to avoid or cope with dosage imbalance can occur either through changes in the non-duplicated partner or through changes in the non-deleterious duplicate. Further, we find selection to correct dosage imbalance can impact the mass and stickiness of some critical interface resides. The combination of these results indicates that the fate and functionality of a duplicate gene and their interacting parters is in part dictated by protein structure.

We also find that dosage imbalance can destabilize a protein, effectively pushing it into a fitness valley. Once dosage balance is restored, this gene is then free to explore mutational space like any other duplicate, but starting from a different starting position on the fitness landscape. From our simulations, we find that once dosage balance had been restored, the duplicate is able to recover some of its stability by climbing up a different fitness peak. Though this fitness peak is suboptimal, the fact that a novel area of mutational space is explored is notable. The exploration of distance mutational space suggests that dosage balance may not only act as a transition state to subsequent neo- or sub-functionalization (Teufel et al., 2016), but actually promote the appearance of these functional changes. In fact, other studies have noted that the evolution of promiscuity, an important step towards functional change, is often due to protein-destabilizing mutations, with additional refinement of novel functionality leading to structural restabilization (Dellus-Gur et al., 2015; Petrie et al., 2018; Sikosek and Chan, 2014; Tokuriki and Tawfik, 2009)

Further, we find that how interacting partners cope with dosage imbalance is stochastic in nature, and we introduce several terms to describe how dosage imbalance affects the evolution of interacting partners. The stochasticity of our experiments suggests that mechanisms not traditionally considered may describe how duplicated genes diverge and how networks of interacting proteins cope with dosage imbalance. For example, the loss of functionality, such as the ability to bind, is often attributed to deleterious mutations in a duplicate gene. While many duplicate genes may in fact lose functionality this way, our results imply that loss of function can also be achieved by, or result in changes to, the original interaction. Additionally, the terms introduced here, *defunctionalization, isofunctionalization*, and *refunctionalization*, to categorize the ways in which a duplicated gene’s interacting partners respond or adapt to the duplication, offer a conceptual framework for describing the consequences of gene duplication in a larger interaction network.

We would like to mention that the phrase isofunctionalization is often used in a different context to refer to non-homologous enzyme isoforms (Omelchenko et al., 2010). Here we use the term in a congruent fashion to denote the loss of ancestral function while maintaining extant function in the context of protein evolution. The underlying idea behind the concept remains the same; simply, a set of proteins or enzymes perform an equivalent function, they just differ in how they do it. The concept of refunctionalization has also been previously introduced, though it has been used as blanket term for a functional change (Beerhues and Liu, 2009; Vecchi, 2012). This concept does differ from our use of the term, which we use to describe the temporary loss of function in evolutionary time.

While the prevalence of any of these mechanism across genome evolution is unknown, our findings suggest that the process of duplicate gene divergence may be more complex than previously appreciated. Further, our results demonstrate that the presence of a duplicated gene can shape how duplicated and non-duplicated interacting partners evolve, independent of the fate of the redundant gene. In fact, the duplicated gene may ultimately be lost over the course of evolution, but its presence may have a lasting impact on the evolution of other members in its interaction network.

Granted, these hypotheses of how functional changes occur in duplicates are based solely on simulations. The prevalence of remodeling at the protein-protein interface, to cope with dosage imbalance, in naturally evolving genomes is unknown. Our observations could be a special case associated with smaller proteins with one interacting partner. Exploring the evolution of more complex protein networks, where either the entire network or substantial portions of interacting partners are duplicated, would be ideal. However, simulations of just three interacting partners can take weeks to run for a single selection scheme. Simulating a larger system would also require running the simulations for longer, in order to reach mutation-selection balance. Hence, computational constraints limit the scope of this study. Our observations could also be affected by how the evolutionary process was simulated. The accelerated origin-fixation model used here changes the order in which substitutions are accepted across evolutionary time (Teufel and Wilke, 2017). However, this reordering was shown to have only a minor influence when compared to evolutionary experiments which did not use this accelerated model (Teufel and Wilke, 2017). In fact, this model has been shown to generate realistic variation in alignments of protein sequences (Jiang et al., 2018).

Even though our study is based on a simplified protein-interaction network with only three partners, it still generates novel hypotheses about how duplicated genes diverge and how protein interfaces evolve. Further, it seems unlikely that more complex natural systems would display less variation than observed in our small simulated system. Our study suggests that, even with just three interacting proteins, a wealth of different evolutionary pathways are possible. Additionally, our results demonstrate that duplicated proteins can have long lasting effects on how interacting partners evolve, and these effects are a function of both stochastic events and protein structure. In total, our findings suggest a structurally-aware and network-wide perspective is essential to understanding the many fates and consequences of gene duplication.

## Methods

We construct a simulation of protein evolution with the use of an accelerated origin-fixation model (Teufel and Wilke, 2017). The simulation is initialized with a small ubiquitin-like protein complexed with a peptide it binds as the resident genotype (PDB: 2EKE, Duda et al. 2007). This protein complex has two subunits, which we refer to as A and B. At each step in the simulation, a novel genotype is created by mutating a single amino acid to a non-resident amino acid. The mutated protein is then locally repacked 5 Å around the mutation. The stability of each subunit (Δ*G*_subunit_) and the stability of binding (Δ*G*_binding_) are evaluated with Rosetta’s all-atom score function (Leaver-Fay et al., 2011; Rohl et al., 2004).

To convert protein stability and binding into fitness, we use a soft-threshold model. This model assumes that the protein’s fitness is given by the fraction of proteins in the ground state in thermodynamic equilibrium (Chen and Shakhnovich, 2009; Serohijos et al., 2012; Wylie and Shakhnovich, 2011). This assumption results in a sigmoidal fitness function (specifically, the Fermi function), where very stable proteins have a fitness of one and fitness declines as stability passes through a threshold value. We calculate the fitness of contributions of stability and binding as

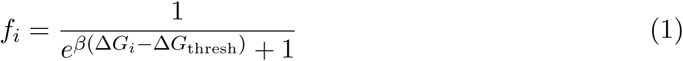
where *β* is the inverse temperature, Δ*G_i_* is the structural stability or stability of the binding interface for protein *i*, and Δ*G*_thresh_ is the threshold at which the protein has lost 50% of its activity. To combine the fitness contributions of binding and stability into a single fitness measure, we log-transform fitness as

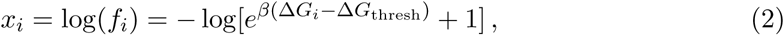
and sum over the fitness contributions of binding and stability (Kachroo et al., 2015). When penalizing binding, we use subtraction rather than addition of the fitness contribution of binding. Using this metric of fitness allow us to express the probability that the mutant genotype will replace the resident genotype as

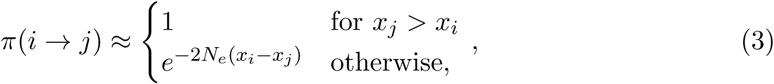
where *N_e_* is the effective population size (Teufel and Wilke, 2017).

At each step in the simulation, the probability of replacement of the resident genotype with a mutant genotype is evaluated with Eq. 3. Our simulations assume that *β* = 1, each Δ*G*_thresh_ component is set to half of the initial stability or stability of binding, and *N_e_* = 1. A burn-in phase is run for 1000 substitutions to ensure that steady-state sampling behavior is being exhibited.

To quantify the functionality of binding we set a functionality threshold (Δ*G*_functional_), and we consider Δ*G* of binding in excess of this threshold as non-functional. We derive the threshold value by setting the left-hand side of Eq. 3 equal to 10^−4^ and solving for Δ*G_i_*. Assuming the state of the system is at Δ*G*_thresh_, we find

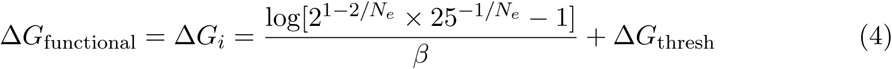

Seven different selection mechanisms are implemented, and we simulate 100 replicates of each of these conditions for 2000 substitutions each. These simulations all assume that selection acts on the stability of each subunit and they differ in how selection pressure on binding is imposed. Fig. 1 illustrates each of the selection schemes we simulate. Two of these simulations are controls and do not include a duplication event (selection schemes 1 and 2). To examine how the duplication of a subunit affects evolutionary dynamics, we carry out five more sets of simulations (selection schemes 3-7). In these simulations, we assume that the B subunit is duplicated, and we refer to the duplicate protein as B’. A subset of these experiments are also repeated assuming that A is the duplicated protein (so that we have A and A’) rather then B. We run simulations were A’ is the duplicated subunit for selection schemes 3, 5, and 6. We also simulate the evolution of an antifungal protein (PDB: 4GVB, Allen et al. 2013) under selection schemes 3, 5, and 6 to examine if our findings are specific to a particular protein structure. *G*_functional_ and the Δ*G*_thresh_ values are also recalculated for this system.

When analyzing how functional change occurs under each of these different selection scenarios, we compare the location of substitutions, both in terms of which subunits they occur in and their position relative to the binding interface, between the bind both (selection scheme 3) and bind B and not B’ (selection scheme 6) experiments. We choose these two experiments for comparison because they both include five terms in their fitness functions. This choice is made because the inclusion of other scenarios with fewer terms in the fitness function can affect the point of mutation-selection balance. Interface residues are considered to be those within 8 Á of the binding interface. The software package SPIDDER (Porollo and Meller, 2007) is used to determine these residues from the structure used to initialize each of our experiments.

## Code

The software, results, and analysis tools used for these experiments will be archived on github.

## Acknowledgements

This work was supported by the National Institutes of Health grant R01 GM088344 to C.O.W. Additional support was provided by the National Science Foundation Cooperative Agreement no. DBI-0939454 (BEACON Center) and grants to E.M.M. from the National Institutes of Health, National Science Foundation, and Welch Foundation (F1515).

---

**Table 1:** Definitions of functionalization pathways. Each pathway is defined based on the ability a subunit has to bind its current and/or ancestral binding partner.

| Functionalization type | Bind current partner | Bind ancestral partner |
| --- | --- | --- |
| Defunctionalization | Lost and never regained | N/A |
| Isofunctionalization | Never lost | Lost at any point |
| Refunctionalization:<br>hard isofunctionalization | Lost and later regained | Permanently lost |
| Refunctionalization:<br>soft isofunctionalization | Lost and later regained | Intermittently lost |
| Refunctionalization:<br>reacquisition | Lost and later regained | Lost and later<br>permanently regained |

